# Multimodal Behavior Scoring quantifies depression-like severity across chronic stress models and identifies stress-resilient mice

**DOI:** 10.64898/2026.01.26.701905

**Authors:** Yue Tu, Qinghui Fu, Yixin Li, Canghao Sun, Yanan Zhu, Jing Deng, Haoning Qin, Xianlin Zeng, Yuchuan Wang, Shulan Qiu, Weixiong Zhang

## Abstract

**Introduction:** Rodent chronic-stress models are central to preclinical depression research, yet behavioral phenotypes are often assessed assay-by-assay, which can reduce cross-cohort comparability and mask heterogeneity across symptom domains (e.g., anhedonia, anxiety-like behavior, and behavioral despair). An integrative behavioral index may provide a more robust and interpretable measure of overall stress-induced severity and facilitate stratification of susceptible versus resilient individuals.

**Methods:** We developed the Multimodal Behavior Scoring (MBS) algorithm to integrate outcomes from standard assays—sucrose preference test (SPT), open field test (OFT), and forced swim test (FST)—into a unified severity score. MBS was evaluated in two independent paradigms, chronic unpredictable mild stress (CUMS) and chronic social defeat stress (CSDS), quantifying group discrimination, reproducibility (intraclass correlation coefficient, ICC), and phenotypic stratification.

**Results:** MBS robustly separated stressed from control mice (CUMS: +39.8%, p<0.0001; CSDS: +121.8%, p<0.0001) and showed high cross-cohort reproducibility (ICC>0.88). MBS further identified stress-resilient subpopulations and uncovered paradigm-specific behavioral signatures, with despair-dominant patterns in CUMS (FST–MBS, r=0.61) and anxiety-centric patterns in CSDS (OFT–MBS, r=-0.74). Excluding TST did not reduce discrimination performance.

**Conclusion:** MBS provides an integrative framework for quantifying depression-like behavioral severity and heterogeneity, enabling streamlined protocols and improving the translational utility of preclinical cohorts for biomarker discovery and antidepressant screening.

## 1. Introduction

Major depressive disorder (MDD) is a debilitating global health concern, affecting approximately 5% of adults worldwide and representing a leading cause of disability [1, 2]. Despite decades of research, progress in developing novel antidepressants has stalled, with numerous clinical trials failing despite promising preclinical findings [3, 4]. A major translational obstacle is the limited capacity of animal models to capture the multifaceted human depressive syndrome and the lack of objective, integrative quantification.

Rodent models, particularly mice subjected to chronic stress paradigms, are widely used in depression research. These models recapitulate core behavioral endophenotypes of depression—including anhedonia, despair, and social withdrawal—and share conserved underlying neurobiological mechanisms involving monoaminergic system disturbances, hypothalamic-pituitary-adrenal (HPA) axis dysregulation, and neuroplasticity deficits [5-8]. Among these, the Chronic Unpredictable Mild Stress (CUMS) paradigm induces depression-like states via varied, low-intensity stressors [9, 10], whereas the Chronic Social Defeat Stress (CSDS) model effectively simulates the profound impact of psychosocial trauma [11, 12]. Both models support mechanistic studies, therapeutic screening, and the study of resilience among stress-exposed individuals.

Currently, depression-like behaviors in these models are quantified primarily through established behavioral assays: the Sucrose Preference Test (SPT) [13] measures anhedonia, the Open Field Test (OFT) [14] evaluates anxiety and locomotor activity, and the Forced Swim Test (FST) [15] and the Tail Suspension Test (TST) [16] assess behavioral despair [17]. Despite their widespread use, these assessments face significant limitations. Critically, depression manifests as a heterogeneous array of symptoms [18], yet conventional assays evaluate these modalities in isolation (e.g., SPT for anhedonia, FST for despair), lacking a comprehensive integrative measure of overall severity. This underscores the need for integrated, quantitative metrics capturing overall severity [19]. Additionally, manual scoring introduces subjectivity and inter-rater variability [20], while inherent behavioral noise and low sensitivity often obscure subtle phenotypic differences, particularly in mild cases or during recovery phases [21, 22]. Furthermore, existing metrics are inadequate in predicting individual susceptibility, resilience, or longitudinal disease progression [23-25]. reliable identification of stress-resistant individuals within exposed cohorts remains challenging but essential [26, 27]. Variability in test performance and interpretation across different stress models [28, 29] also raises concerns regarding cross-model validity and reproducibility. The lack of standardized, objective, and integrative assessment contributes significantly to reproducibility issues across laboratories or even experiments [30].

To address these issues, we developed Multimodal Behavior Scoring (MBS), a robust algorithm that integrates key metrics from established behavioral tests into a single continuous severity score, enabling a more holistic and objective assessment of depression-like behavior. The framework is flexible, allowing inclusion or exclusion of individual modalities (such as the TST) depending on experimental objectives. We validated MBS in both CUMS and CSDS models, evaluating cross-model reliability and generalizability, and demonstrated its utility for tracking progression, recovery, and recurrence after stress exposure and cessation. Using MBS, we identified stress-resistant mice that maintain normal behavioral function despite chronic stress exposure, enabling investigation of resilience mechanisms. Notably, our results also questioned the necessity of the TST in core behavioral batteries, suggesting minimal or counterproductive contribution relative to the integrated SPT-OFT-FST triad.

Collectively, we present a validated computational approach for integrating multimodal behavioral data into a unified quantitative measure of depression severity. By addressing fragmented and subjective assessments, MBS provides a standardized tool to enhance reproducibility, facilitate identification of stress-resistant phenotypes, and help bridge translational gaps in depression research.

## 2. Materials and Methods

### 2.1 Animals

C57BL/6J male mice (4–6 weeks, 18–22 g) were obtained from the Centralized Animal Facility (CAF) of The Hong Kong Polytechnic University (PolyU). All animal procedures were approved by the Animal Subjects Ethics Sub-committee of PolyU (No. 23-24/889-HTI-R-STG, No. 24-25/1083-HTI-R-STG) and conducted in accordance with the guidelines of the Department of Health of the Hong Kong SAR.

Mice were housed in barrier facilities under controlled conditions (22 ± 1 °C; 60 ± 10% humidity) with ad libitum access to food and water, and maintained on a 12:12-hour light/dark cycle (lights on 8:00–20:00) unless stated otherwise.

### 2.2 Study Design and MBS Workflow

We employed two complementary yet mechanistically distinct stress paradigms: chronic unpredictable mild stress (CUMS), which models chronic environmental adversity through randomized low-intensity stressors, and chronic social defeat stress (CSDS), which models psychosocial trauma via resident–intruder confrontations. These paradigms capture different aspects of depressive etiologies and together provide a more comprehensive evaluation of depressive phenotypes, enabling delineation of shared and model-specific vulnerability markers and supporting characterization of stress-resistant phenotypes (Figure 1A).

**Figure 1.**
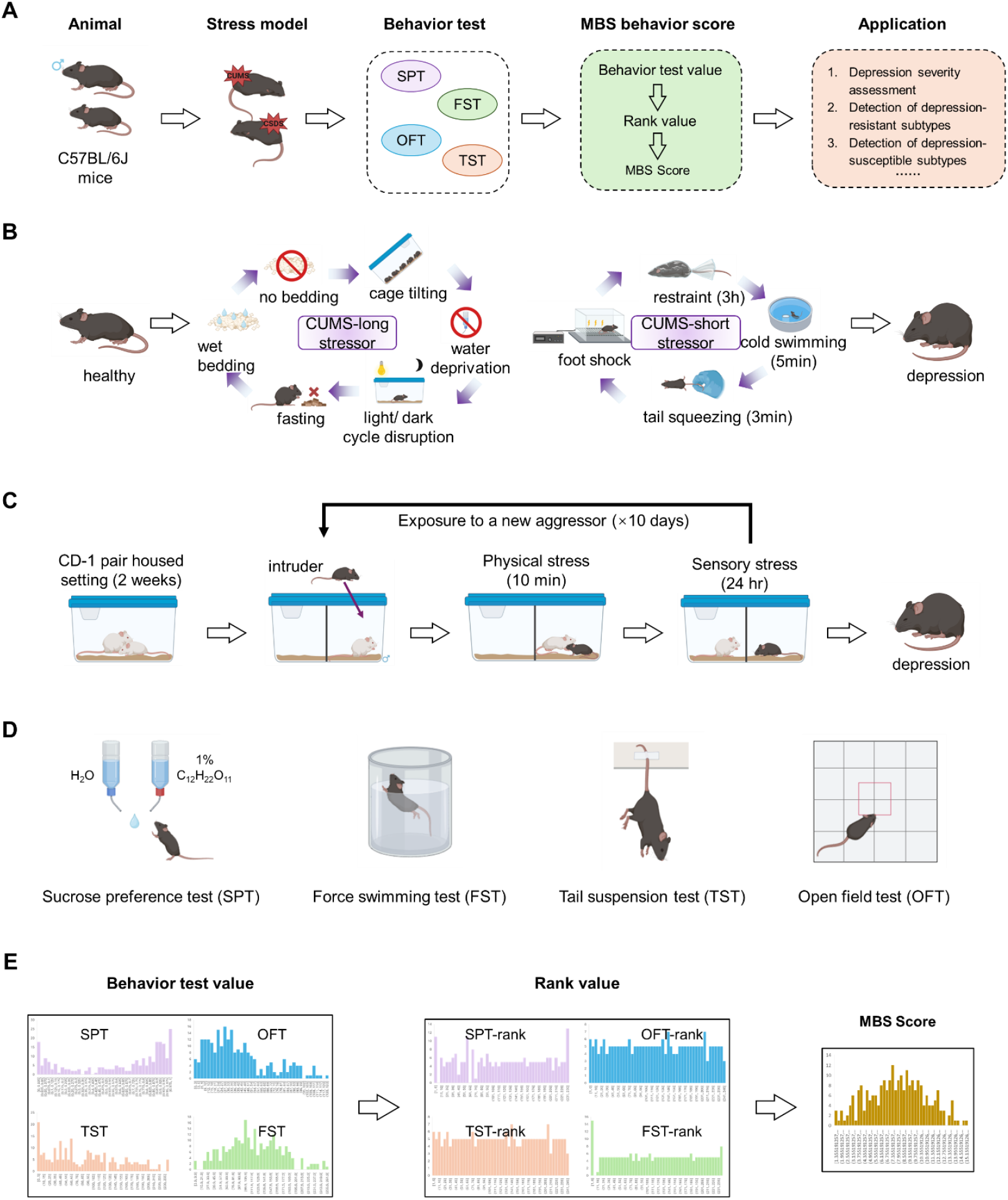
Overview of the methodological framework. (**A**) Flowchart depicting animal experiments and behavioral assessment procedures. (**B**) Chronic Unpredictable Mild Stress (CUMS) model. (**C**) Chronic Social Defeat Stress (CSDS) model. (**D**) Schematic representation of behavioral tests. (**E**) Principle of the Multimodal Behavior Scoring (MBS) algorithm.

The overall experimental design established a unified pipeline for multimodal behavioral phenotyping. CUMS (Figure 1B) and CSDS (Figure 1C) cohorts underwent a standardized behavioral battery (Figure 1D), and behavioral readouts were integrated by the Multimodal Behavior Scoring (MBS) algorithm to generate a continuous severity score and support subsequent validation and phenotypic stratification. The computational architecture of MBS (Figure 1E) applies cohort-specific nonparametric rank transformation followed by linear integration of multimodal inputs, providing an analytical foundation for downstream analyses, including identification of stress-resilient and stress-susceptible subtypes and longitudinal tracking following stress cessation.

### 2.3 Animal Models of Depressive-like Behavior

#### Chronic Unpredictable Mild Stress (CUMS)

To induce depression-like behaviors, mice were subjected to a 4-week CUMS protocol with minor adaptations [31-33]. Each day, animals received one long-duration and one short-duration stressor, randomly selected from a predefined list to ensure unpredictability. Long-duration stressors included 24 h food or water deprivation, 16 h cage tilting (45°), 16 h strobe light exposure, 24 h housing without bedding, and 24 h housing with wet bedding. Short-duration stressors included 5 min cold water swim (4°C), 3 min tail pinch, 2 h restraint stress, and 10 min foot shock. Stressors varied daily in type and timing to prevent habituation and simulate chronic psychosocial stress.

#### Chronic Social Defeat Stress (CSDS)

The CSDS model was performed for 10 consecutive days using a standard resident–intruder paradigm. Male CD-1 mice (4–6 months old; Laboratory Animal Service Centre, The Chinese University of Hong Kong) served as resident aggressors, and male C57BL/6J mice (6–7 weeks old) served as intruders. Aggressors were screened 3 days before CSDS to select mice with adequate aggressiveness.

One day before the first defeat session, selected aggressors were acclimated overnight in one side of a perforated, divided cage. Each day, a novel C57BL/6J intruder was introduced into the aggressor compartment for a 5–10 min defeat episode, then moved to the opposite compartment to allow continuous sensory contact through the divider for the next 24 h. A new aggressor was used each day.

Sessions were monitored to ensure consistent aggression (≥1 aggressive bout/min; each bout 5–10 s). Aggressors displaying affiliative behaviors (e.g., grooming) were excluded and replaced. Control mice were housed in identical divided cages with another control mouse but without physical interaction.

### 2.4 Behavioral testing procedures

To assess depression- and anxiety-like behaviors following chronic stress, mice underwent the sucrose preference test (SPT), open field test (OFT), forced swim test (FST), tail suspension test (TST), and the social interaction test (SIT; CSDS only). Except for SPT, all assays were conducted under dim light in a quiet testing room after 1–2 h habituation. Sessions were video-recorded and analyzed using Any-Maze (Stoelting, IL, USA) unless otherwise specified.

SPT was performed as previously described [34, 35] to assess anhedonia. Mice were habituated to two bottles of 1% sucrose for 24 h, followed by two bottles of water for 24 h, with bottle positions switched every 12 h. After 24 h food and water deprivation, mice were given one bottle of water and one bottle of 1% sucrose for 2 h starting at 20:00, with positions exchanged after 1 h. Sucrose preference was calculated as sucrose intake/total fluid intake.

OFT assessed locomotion and exploration in a 40 × 40 × 40 cm opaque chamber. Mice were placed in a corner and recorded for 6 min, excluding the first minute from analysis. The arena was cleaned with 75% ethanol between trials. Time spent in the center and periphery was quantified.

FST assessed behavioral despair in a transparent cylinder (15 cm diameter, 30 cm height) filled with 23–25°C water to 18 cm depth. Mice were tested for 6 min and immobility time was quantified; animals were dried before returning to home cages.

TST assessed despair-like behavior by suspending mice via tail tape (∼1 cm from the tip) with a rubber tube to prevent climbing. Mice were suspended 20 cm above the floor for 6 min, and immobility during the last 4 min was analyzed.

For CSDS mice, SIT was conducted 24 h after the final defeat to assess social avoidance. Mice were tested in a 44 × 44 × 38 cm arena containing a wire-mesh enclosure (10 × 6.5 × 38 cm) across two 2.5-min trials (target absent, then target present with a novel CD-1), separated by 30 s. Time in the interaction zone was recorded, and the social interaction ratio was calculated as target-present/target-absent interaction time.

All behavioral testing and analyses were conducted blinded. Sufficient recovery time was allowed between assays, and test order was counterbalanced across groups to minimize sequence effects.

### 2.5 Multimodal Behavior Scoring (MBS)

To quantitatively integrate heterogeneous behavioral metrics and overcome the modal fragmentation inherent to classical behavioral tests, we developed the Multimodal Behavior Scoring (MBS). MBS is a deterministic algorithm that projects individual-level behavioral data into a unified latent severity space, designed to capture depressive load across multiple modalities. The method operates via three sequential stages: metric transformation, rank space embedding, and normalized integrative projection.

Let 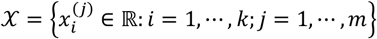 denote the raw behavioural data matrix, where *k* is the number of animals and *m* the number of behavioural features (e.g., SPT, OFT, FST, TST). Each variable 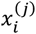 represents the observed behavioral outcome for mouse *i* in test *j*. These features differ in range, directionality, and distribution, rendering them non-comparable in raw form.

We define a polarity function *π*: {1, ⋯, *m*} → {−1,1} such that:

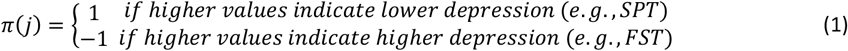

Each variable is transformed into a relative rank space via:

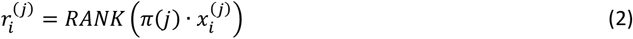

where *RANK*(·) denotes the position of the value in the sorted list of transformed scores within modality *j*, with rank 1 assigned to the most severe behavioral phenotype.

Let 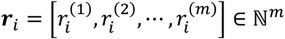 be the rank vector of mouse *i*.

#### Definition (MBS Score)

The raw score for subject *i* is completed as the mean normalized rank across selected modalities:

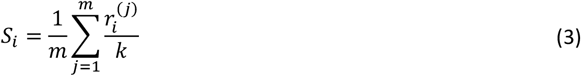

To normalize this to a bounded range and enhance comparability across experiments, we define the final MBS score as:

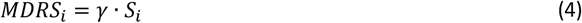

where *γ* > 0 is a tunable scaling constant (we set *γ* = 16 in practice), such that 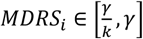.

This mapping ensures interpretability and comparability across experimental batches, regardless of cohort size or test modality. The MBS construction exhibits several desirable theoretical properties that support its use as a core metric in preclinical behavioral neuroscience.

The MBS framework, though algorithmically simple, possesses a set of well-defined mathematical properties that support its theoretical validity and practical robustness. These properties ensure the stability, interpretability, and extensibility of the score across varying behavioral inputs and experimental conditions. Below, we formally characterize several key invariance and distributional features of MBS.

#### Lemma 1 (Permutation Invariance)

Let *σ*: {1, ⋯, *k*} → {1, ⋯, *k*} be any permutation of the animal index. Then, the distribution of {*MDRS*_*i*_} is invariant under *σ*. That is, MBS scores depend solely on relative behavioural positioning, not animal identity.

#### Lemma 2 (Scale Invariance)

Let behavioural feature *j* be subject to an affine transformation 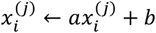 for all *i*, with *a* > 0. Then, the MBS scores remain unchanged. Thus, MBS is robust to unit scaling and shifts in measurement (e.g., converting immobility time from seconds to z-scores).

#### Proposition 1 (Asymptotic Normality)

Let the entries 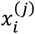 be independently and continuously distributed. Then, by the Lyapunov central limit theorem, the distribution of *MDRS*_*i*_ converges to a Gaussian as *m* → ∞ and/or *k* → ∞, facilitating parametric statistical inference.

#### Remark (Modality Modularity)

MBS is explicitly modular in the choice of behavioral modalities *m*, allowing for the dynamic inclusion or exclusion of specific tests. For instance, a 3D variant (SPT–OFT–FST) can be directly compared to a 4D variant (including TST) by rescaling via *γ* and using shared rank embedding.

This rank-based formulation provides robustness to outliers, skewed distributions, and inter-test heteroskedasticity—issues that commonly undermine raw-score-based composite indices. Furthermore, since all features are rank-normalized, their contribution to the final MBS score is equal by construction, regardless of their absolute variance or dynamic range.

In practical application across both CUMS and CSDS models, we used MBS to calculate depressive severity scores from behavioral data collected from standard tests. The resulting MBS distributions were approximately Gaussian, and animals with extreme scores could be reliably identified as highly stress-susceptible or stress-resilient. Notably, MBS allowed us to detect sub-threshold or compensatory behavioral patterns that were not apparent in any individual test, offering a more nuanced phenotype stratification than categorical classification or test-by-test significance testing.

By embedding multimodal behavioral readouts into a uniform, interpretable space, MBS provides a scalable and theory-grounded alternative to conventional behavioral indices. It not only improves statistical power in the face of noisy and heterogeneous data but also facilitates the phenotyping of intermediate states, the detection of resilient individuals, and the reproducible tracking of behavioral recovery across models and time points.

### 2.6 Statistical analysis

All analyses were performed in GraphPad Prism 9.0. Data are presented as mean ± SEM unless otherwise stated. Two-group comparisons (e.g., control vs. stress) were conducted using unpaired Welch’s t-tests. Normality was assessed using Shapiro–Wilk tests; nonparametric tests were considered but not required. Significance thresholds were *p* < 0.05 (*), *p* < 0.01 (**), *p* < 0.001 (***), and *p* < 0.0001 (****). Multiple-comparison correction was not applied unless specified, as analyses focused on preplanned group contrasts. Associations between MBS and individual behavioral measures were evaluated using Pearson’s correlation (r). No animals or data points were excluded unless predefined artifact criteria (e.g., incomplete trials or equipment error) were met.

## 3. Results

Building on the experimental design and MBS workflow (Figure 1), we evaluated the robustness and interpretability of MBS across independent CUMS and CSDS cohorts. We first tested whether rank transformation improves cross-cohort comparability (Section 3.1), then assessed MBS sensitivity and cross-model consistency in quantifying depression-like severity (Section 3.2). We next applied MBS to phenotype stratification to delineate susceptibility and resilience (Section 3.3), examined the contribution of TST to multimodal scoring (Section 3.4), and finally demonstrated longitudinal tracking of recovery and recurrence following stress cessation and re-exposure (Section 3.5).

### 3.1. Rank transformation enhances cross-cohort behavioral comparability

The integration of multimodal behavioral data across cohorts requires robust normalization to mitigate inherent variability. MBS addresses this by applying cohort-specific rank transformation, converting heterogeneous metrics (SPT, OFT, and FST) into uniform percentile distributions. In both CUMS (Figure 2A–C) and CSDS (Supplementary Figure S1A–C), rank-transformed values retained strong monotonic relationships with the original measures (Pearson r > 0.92, p < 0.0001), indicating preservation of phenotypic information during rescaling. Importantly, rank transformation generated stable uniform distributions that neutralized baseline shifts across cohorts, enabling cross-cohort integration within a common metric space.

**Figure 2.**
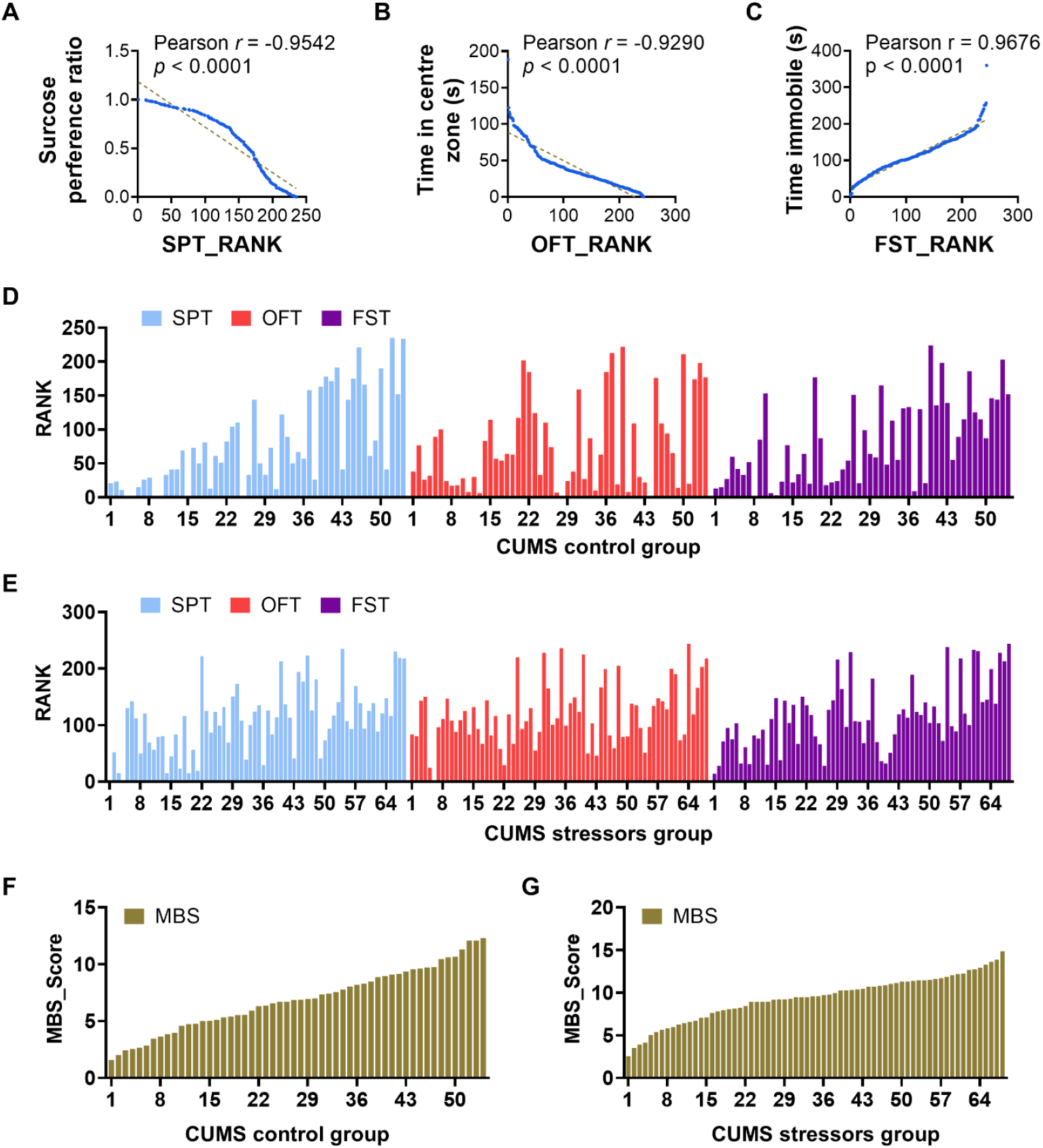
Rank transformation enables robust cross-cohort integration of behavioral data in the CUMS model. (**A**) Correlation between raw and rank-transformed scores in the sucrose preference test (SPT). (**B**) Correlation between raw and rank-transformed scores in the open field test (OFT). (**C**) Correlation between raw and rank-transformed scores in the forced swim test (FST). (**D**) Rank-based trajectories of individual behavioral metrics (SPT, OFT, FST) sorted by ascending MBS scores in CUMS control group. (**E**) Rank-based trajectories of individual behavioral metrics (SPT, OFT, FST) sorted by ascending MBS scores in CUMS stressors group. (**F**) Ordered bar chart of MBS in CUMS control group. (**G**) Ordered bar chart of MBS in CUMS stressors group.

Ordering mice along the MBS severity continuum further revealed close alignment between composite scoring and individual behavioral modalities. When stratified by ascending MBS values (CUMS: Figure 2F/G; CSDS: Supplementary Figure S1F/G), rank-transformed behavioral trajectories (Figure 2D/E; Supplementary Figure S1D/E) showed striking covariation with MBS. Individual-level analyses similarly demonstrated coordinated directional shifts in SPT, OFT, and FST ranks within MBS-defined subgroups (Supplementary Figures S2 and S3), supporting coherent multimodal integration without distorting phenotypic profiles.

Together, these findings establish rank transformation as the key mathematical step that suppresses batch-specific artifacts while preserving neurobehavioral signals, forming the basis for subsequent phenotype discovery and cross-model validation.

### 3.2. MBS quantifies depression severity with high sensitivity and cross-model consistency

The MBS approach demonstrated exceptional discriminative power on CUMS cohorts. Stressed mice exhibited profound depression-like states characterized by a 39.8% increase in MBS scores compared to controls (9.070 ± 2.581 vs 6.489 ± 2.445; p < 0.0001, Figure 3A). This composite severity metric captured coordinated deterioration across core behavioral domains: sucrose preference (SPT) declined by 19.6% (58.47% ± 0.3292 vs 72.73% ± 0.3222 in controls; p = 0.0089, Figure 3B), reflecting established anhedonia phenotypes; exploratory activity in the open field (OFT) diminished by 50.3% (28.49 ± 20.91s center time vs 57.35 ± 37.91s in controls; p < 0.0001, Figure 3C), indicating heightened anxiety; and immobility duration in forced swim tests (FST) increased by 32.8% (130.4 ± 51.74s vs 98.20 ± 50.17s in controls; p = 0.005, Figure 3D), demonstrating behavioral despair. Critically, MBS scores maintained biologically coherent correlations with these core modalities, showing significant inverse associations with reward sensitivity (SPT: r = −0.5898, p < 0.0001, Figure 3E) and exploratory drive (OFT: r = −0.3967, p < 0.0001, Figure 3F), while exhibiting strong positive correlation with behavioral despair (FST: r = 0.6077, p < 0.0001, Figure 3G). This integrated quantification outperformed isolated behavioral results in phenotypic resolution. Our CUMS mice were derived from two cohorts, with the individual results of MBS, SPT, OFT, and FST for each cohort shown in Figure S4.

**Figure 3.**
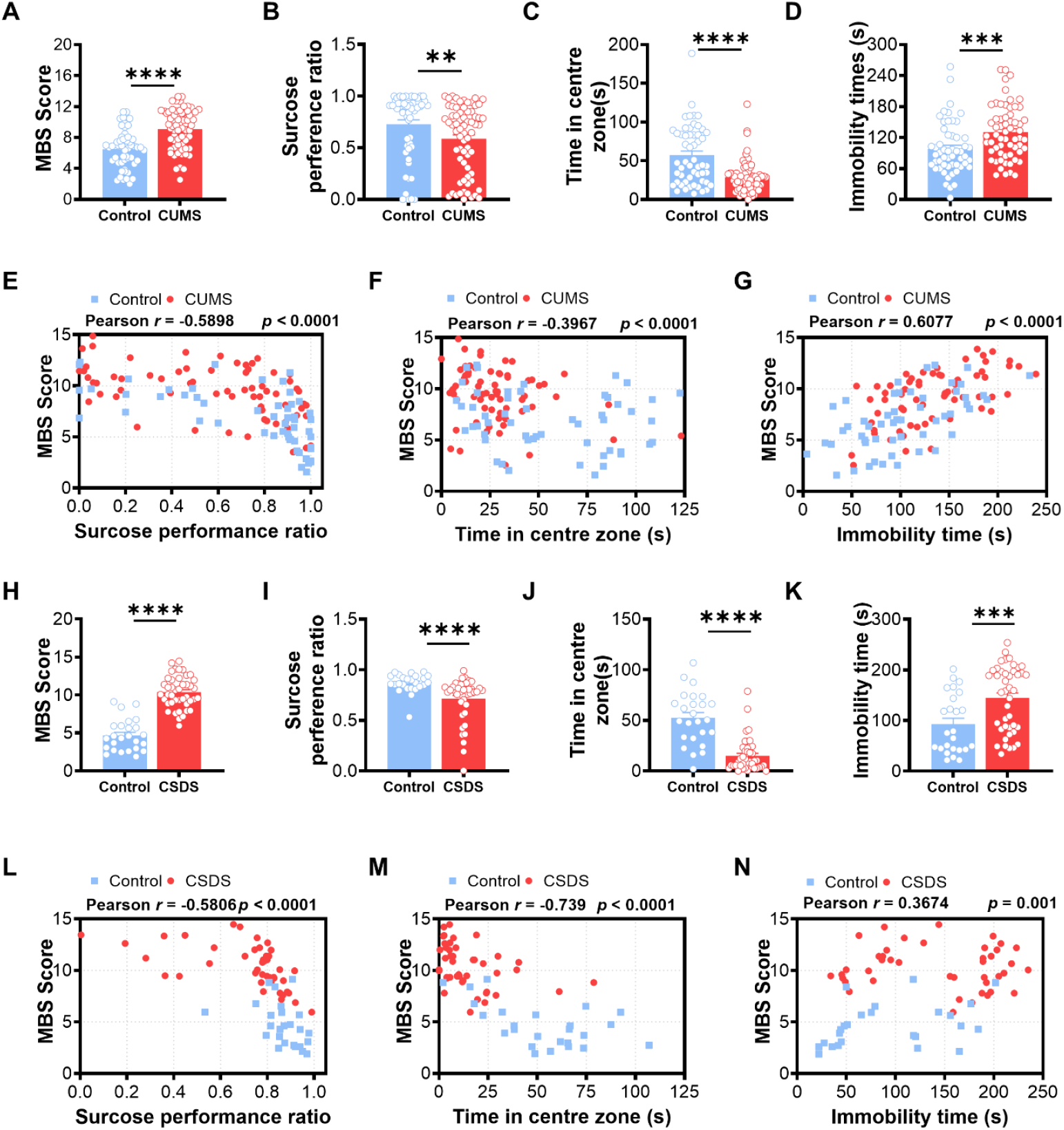
MBS quantifies depression severity with high sensitivity and stress model-specific behavioral integration. (**A**) Elevated MBS scores in CUMS-stressed mice indicate pronounced depression-like states. (**B**) reduced SPT in CUMS model reflects anhedonia. (**C**) Decreased center exploration time in the OFT indicates heightened anxiety in CUMS mice. (**D**) Increased immobility time in the FST suggests enhanced behavioral despair in CUMS mice. (**E**) Negative correlation between MBS scores and sucrose preference (SPT) in the CUMS model. (**F**) Negative correlation between MBS scores and center time (OFT) in the CUMS model. (**G**) Positive correlation between MBS scores and immobility duration (FST) in the CUMS model. (**H**) Significantly elevated MBS scores in CSDS-stressed mice reflect robust depressive phenotypes. (**I**) Decreased sucrose preference in the CSDS model indicates anhedonia. (**J**) reduced OFT center exploration in CSDS mice reflects severe anxiety-like behavior. (**K**) Increased FST immobility duration in the CSDS model reflects moderate behavioral despair. (**L**) Negative correlation between MBS scores and sucrose preference (SPT) in the CSDS model. (**M**) Strong negative correlation between MBS scores and OFT performance in the CSDS model. (**N**) Positive but attenuated correlation between MBS scores and FST immobility in the CSDS model.

On the etiologically distinct CSDS model, MBS similarly detected robust depressive phenotypes while revealing unique pathophysiological features. Mice subjected to chronic social defeat developed severe symptoms reflected in significantly elevated MBS scores (10.39 ± 2.086 vs 4.684 ± 2.079 in controls; p < 0.0001, Figure 3H). Detailed behavioral decomposition uncovered a distinct expression profile: sucrose consumption decreased to 71.41% ± 0.2114, representing a 17.7% reduction versus controls (p < 0.0001, Figure 3I); anxiety-related behaviors manifested as a 71.4% decline in OFT center exploration time (15.04 ± 16.30s vs 52.52 ± 25.86s in controls; p < 0.0001, Figure 3J); whereas behavioral despair exhibited comparatively moderate intensification with FST immobility increasing 55.8% (144.7 ± 64.52s vs 92.85 ± 58.32s in controls; p = 0.0006, Figure 3K). The correlation architecture of CSDS models diverged significantly from CUMS: social stress manifested stronger integration of anxiety-related behaviors (OFT: r = −0.7397, p < 0.0001, Figure 3M), whereas unpredictable stress emphasized behavioral despair (FST: r = 0.6077 in CUMS vs r = 0.3674 in CSDS, Figure 3G vs 3N). Similarly, the individual results for each cohort of CSDS-treated mice are presented in Figure S5.

Cross-model comparative analysis revealed fundamental principles governing depression-like behavior expression across stress models. While both models exhibited conserved anhedonia pathology (SPT-MBS correlations: r = −0.5898 in CUMS vs r = −0.5806 in CSDS; |Δr| < 0.01), their phenotypic architectures diverged in stressor-specific manifestations: CUMS produced despair-dominant phenotypes characterized by stronger FST-MBS associations (r = 0.6077 vs r = 0.3674 in CSDS; |Δr| = 0.2403), whereas CSDS generated anxiety-centered pathophysiology showing enhanced OFT-MBS integration (r = −0.7397 vs r = −0.3967 in CUMS; |Δr| = 0.3430). This differential sensitivity profile demonstrates MBS’s capacity for intrinsic model adaptation— automatically amplifying contributions from dominant behavioral modalities without manual parameter adjustment—a critical feature for modeling heterogeneous psychiatric conditions.

Robust reproducibility was confirmed through four independent biological replicates (Supplementary Figures 2-3, encompassing two CUMS cohorts and two CSDS cohorts). MBS maintained exceptional measurement consistency across cohorts, with inter-replicate intraclass correlation coefficients (CUMS: ICC = 0.88; CSDS: ICC = 0.93) substantially exceeding established reliability thresholds for preclinical research (ICC >0.80 [36]). Crucially, the distinctive correlation topology characterizing each stress model remained stable across cohorts (|Δr| < 0.12 for equivalent modalities), confirming MBS’s ability to preserve pathophysiological signatures despite experimental batch effects.

### 3.3 Phenotypic stratification delineates susceptibility and resilience – Finding stress resilient mice

To integratively assess depressive behaviors and identify stress-resistant phenotypes, we established phenotypic stratification criteria based on multimodal behavioral profiles. Control mice (CON) were defined as unstressed animals with consistently low MBS scores (<7). Within stress-exposed cohorts, resilient mice (RES) maintained control-like performance with MBS <7, whereas susceptible mice (SUS) showed maladaptive responses with elevated MBS scores (>10). These thresholds were derived from cluster analysis of integrated behavioral trajectories, optimized to discriminate phenotypic categories while remaining consistent with established depressive endophenotypes.

CUMS exposure produced heterogeneous responses across domains (Figure 4A–D). resilient mice were indistinguishable from controls across core measures (MBS: RES5.230 ± 1.315 vs CON 4.878 ± 1.642, p > 0.05; SPT: p > 0.05; FST: p > 0.05), whereas susceptible mice showed markedly increased severity (MBS 11.80 ± 1.140; p < 0.001 vs CON and RES), severe anhedonia (SPT: SUS 41.23% ± 5.3 vs RES79.06% ± 0.2177, p < 0.001), and increased despair (FST: SUS 167.0 ± 54.83 s vs RES92.27 ± 26.70 s, p < 0.001). OFT measures suggested an intermediate anxiety-related profile in resilient mice (OFT: RES35.68 ± 33.84 s vs CON 62.57 ± 38.28 s, p < 0.05).

**Figure 4.**
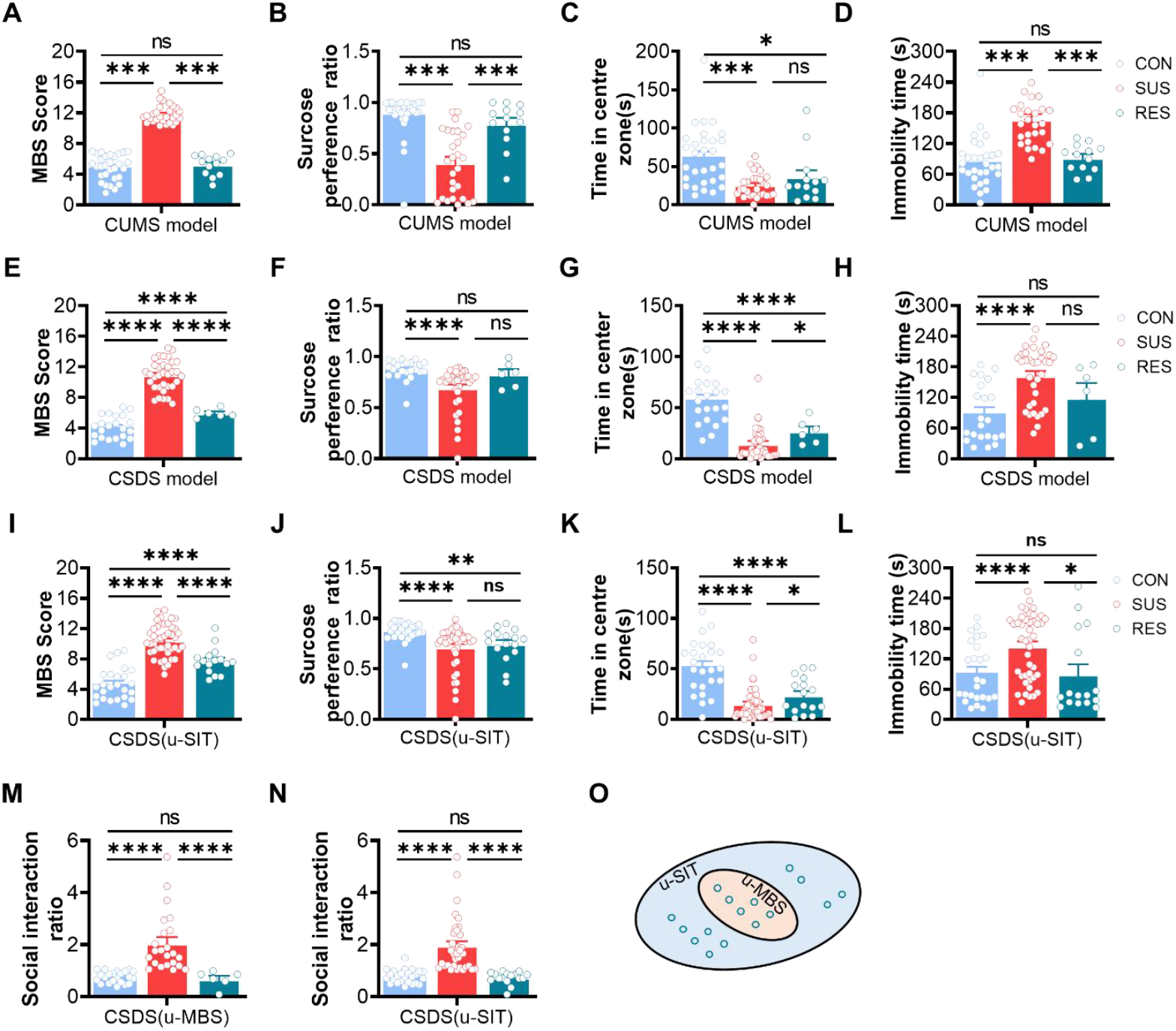
Phenotypic stratification using MBS distinguishes stress-susceptible and resilient individuals across depression models. CUMS model: (**A**) MBS scores differentiate control, resilient, and susceptible mice in the CUMS model. (**B**) SPT is preserved in resilient mice and reduced in susceptible mice. (**C**) OFT center time shows partial anxiety regulation in resilient mice and strong deficits in susceptible mice. (**D**) FST immobility duration is significantly increased in susceptible mice. CSDS model: (**E**) MBS-based stratification identifies control, resilience, and susceptible mice in the CSDS model. (**F**) Resilient mice maintain sucrose preference at control-like levels. (**G**) OFT performance reveals a stepwise gradient of anxiety severity across phenotypes. (**H**) FST immobility is elevated only in susceptible mice, with resilient mice showing intermediate behavior. Validation with SIT: (**I**) SIT ratios of MBS-defined phenotypes show agreement with traditional SIT classifications. (**J**) MBS-defined resilient mice exhibit social behavior like SIT-defined resilient mice. (**K**) SIT-identified resilient mice with high MBS scores show residual behavioral impairments. (**L**) Scatter plot comparing SIT ratios with MBS scores across individuals. Model comparison and refinement: (**M**) Behavioral comparison of mice identified as resilient by utilizing MBS method. (**N**) Behavioral comparison of mice identified as resilient by utilizing SIT method. (**O**) Venn diagram showing overlap between MBS- and SIT-defined resilient subpopulations.

CSDS similarly revealed pronounced stratification (Figure 4E–H). Susceptible mice exhibited elevated MBS (SUS 10.88 ± 1.981) relative to controls (CON 4.124 ± 1.482, p < 0.0001) and resilient mice (RES5.954 ± 0.5867, p < 0.0001). resilient mice showed no significant impairment in sucrose preference, while anxiety-like behavior displayed a graded reduction in OFT center time from CON (57.75 ± 22.65 s) to RES(26.77 ± 11.89 s) to SUS (14.53 ± 16.10 s) (CON–SUS: p < 0.0001; RES–CON: p < 0.0001; SUS–RES: p < 0.05). FST immobility increased significantly only in susceptible mice (SUS 162.1 ± 56.18 s vs CON 88.71 ± 56.45 s, p < 0.0001), whereas resilient mice were intermediate and not significantly different from either group.

Across paradigms, resilience profiles were model-specific: CUMS resilience emphasized preservation of reward and coping-related behaviors with partial retention of anxiety-related vigilance, whereas CSDS resilience was more strongly associated with normalized anxiety responses alongside partially retained passive coping.

Stratification was further validated against the social interaction test (SIT) in CSDS (Figure 4I–L). MBS-defined resilient mice showed comparable social approach to SIT-defined resilient mice (SIT ratio: MBS-RES0.6744 ± 0.3192 vs SIT-RES0.7261 ± 0.2342, p > 0.05). Notably, MBS-RES represented a refined subset of SIT-RES with 37.5% overlap (Figure 4O), and mice classified as resilient only by SIT exhibited residual anxiety-related impairments.

These results demonstrate that the integrative MBS approach robustly distinguishes stress-resilient and susceptible mice, effectively addressing behavioral heterogeneity, and providing a powerful framework for studying resilience mechanisms in depression research.

### 3.4 Empirical assessment reveals a little utility of the tail suspension test

The tail suspension test (TST), conventionally used as an indicator of despair behavior, exhibited minimal discriminative capacity according to our integrated assessment approach. While CUMS-exposed mice showed marginally prolonged immobility compared to controls (89.81 ± 65.12s vs. 70.09 ± 57.37s; p=0.0391, Figure 5A), this difference exhibited the least substantial change among all behavioral measures. Crucially, TST performance exhibited negligible correlation with composite depression severity quantified by MBS (r=0.09849, p=0.2804, Figure 5B), suggesting limited utility as a depression biomarker. This dissociation suggests TST responses capture stress-induced motor inhibition rather than depression-specific pathophysiology.

**Figure 5.**
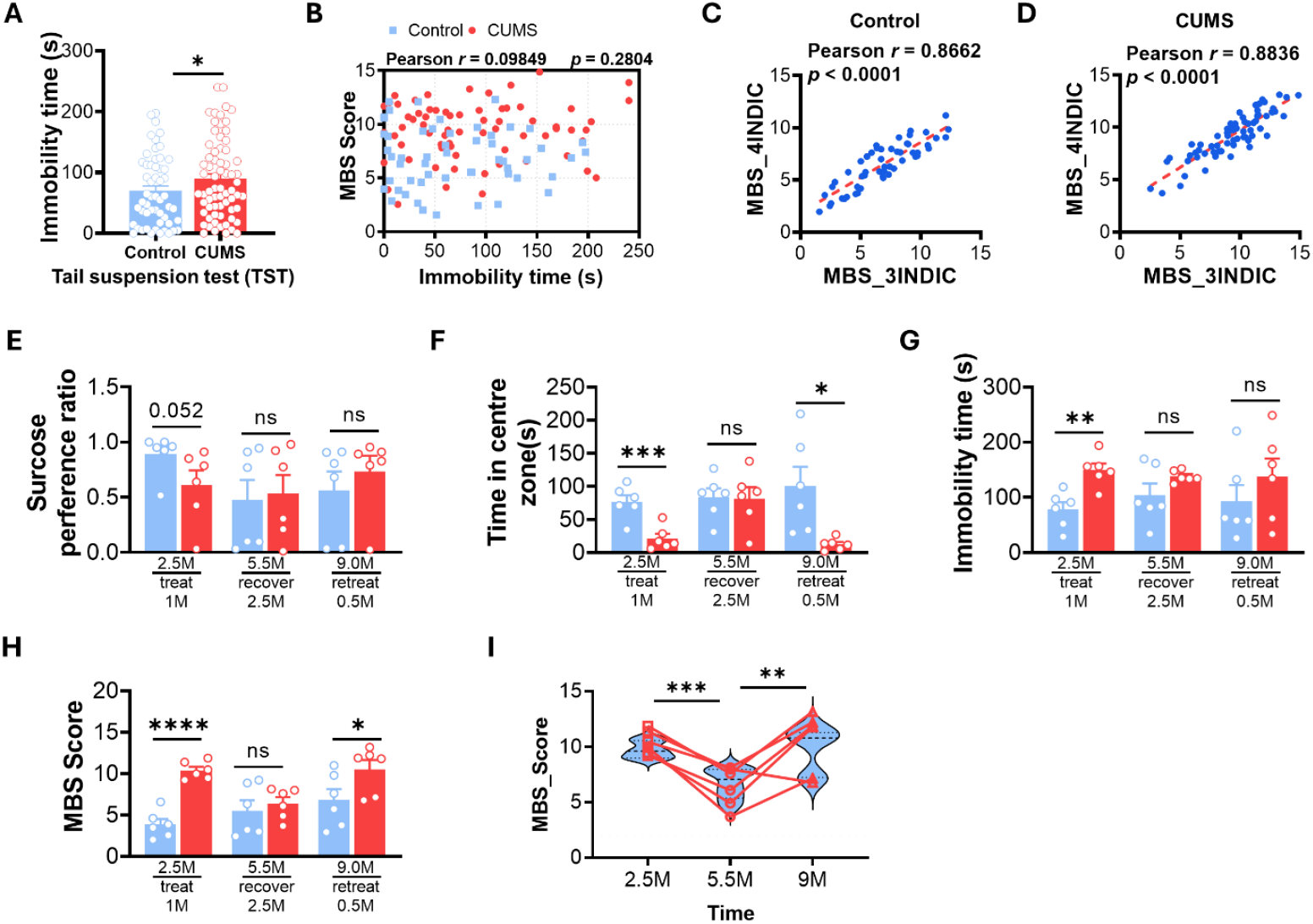
Limited utility of the tail suspension test and dynamic tracking of stress-induced behavioral trajectories via MBS. (**A**) TST immobility duration shows only marginal increase in CUMS-exposed mice. (**B**) TST scores exhibit negligible correlation with composite MBS scores, indicating poor reflection of depressive severity. (**C**) Strong concordance between 4-modal MBS (SPT, OFT, FST, TST) and 3-modal MBS (SPT, OFT, FST) in control mice. (**D**) Strong concordance between 4-modal and 3-modal MBS in CUMS mice, suggesting limited added value of TST. (**E**) SPT of Longitudinal trajectories. (**F**) OFT of Longitudinal trajectories. (**G**) FST of Longitudinal trajectories. (**H**) MBS score of Longitudinal trajectories. (**I**) Longitudinal MBS trajectories of individual mice.

Methodologically, the inclusion of TST contributed little discriminative value to multimodal phenotyping. When comparing the full 4-modal MBS (incorporating TST) against a streamlined 3-modal alternative (SPT/OFT/FST), both control and CUMS cohorts demonstrated essentially identical classification outputs (control: r = 0.8662, p < 0.0001; CUMS: r = 0.8836, p < 0.0001; Figures 5C-D). The less variance between 4-modal MBS and 3-modal MBS confirms that core behavioral domains—anhedonia, exploration deficits, and swim test despair—provide sufficient information for adequate depression phenotyping without TST supplementation.

These results establish that research on depression can achieve equivalent analytical rigor while excluding TST from assessment batteries. This conclusion is consistent with some published results [37]. Such optimization reduces experimental cost and testing duration while minimizing animal stress exposure— critical advances for ethical preclinical research. The persistence of TST usage likely reflects historical convention rather than empirical necessity, an incongruity our data compellingly address through quantitative benchmarking.

### 3.5 MBS dynamics track stress-induced recovery trajectories

Our longitudinal study delineates a dynamic progression of stress-induced depressive phenotypes through sequential induction, recovery, and recurrence phases (Figure 5E-H). When subjected to a 4-week CUMS scheme (T1, age: 2.5 months), mice developed robust depressive-like behaviors evidenced by significantly elevated MBS scores relative to controls (10.39 ± 1.055 vs. 3.838 ± 1.623; p < 0.0001). Following a 2.5-month stress-free recovery period at 5.5 months of age (T2), behavioral deficits were virtually abolished with no statistically distinguishable differences between groups (6.388 ± 1.826 vs. 5.499 ± 3.058; p = 0.2788), indicating phenotypic remission. Then, upon re-exposure to CUMS for a condensed 2-week rechallenge period (T3), previously stressed mice exhibited accelerated symptom recurrence compared to control mice that had not been previously stressed, manifesting as moderately elevated MBS scores that retained statistical significance (10.49 ± 2.783 vs. 6.794 ± 3.244; p = 0.030). This temporal trajectory—characterized by initial susceptibility, full behavioral recovery, and preferential vulnerability to restress—demonstrates the capacity of MBS to detect distinct neuroadaptive transitions.

Serial assessment at critical junctures revealed persistent pathophysiological alterations beneath phenotypic normalization. Longitudinal profiling of individual mice (Figure 5I) uncovered that despite group-level recovery at T2, CUMS subjects maintained elevated recurrence sensitivity during restress. The differential vulnerability was quantified through intragroup comparisons: the T1-to-T2 transition showed a dramatic improvement (p < 0.001), while the T2-to-T3 deterioration occurred with steeper kinetic progression (p = 0.007) relative to primary induction. More profoundly, baseline severity at T1 predicted restress susceptibility at T3 (r=0.8602, p < 0.001), suggesting that MBS captures latent pathophysiology independent of overt symptom expression.

## 4. Discussion

Identifying stress-resistant phenotypes in rodent models is critical for dissecting protective mechanisms against depression [26, 27]. Using MBS, we consistently stratified resilient individuals—mice that maintained relatively normal behavioral function despite chronic stress—thereby providing a quantitative basis for probing resilience-associated molecular and circuit adaptations.

Conceptually, MBS treats depression-like behavior as an integrated, multimodal phenotype rather than a collection of isolated assay readouts. By linearly integrating rank-transformed SPT, OFT, and FST metrics, MBS captures coordinated dysregulation across reward, affective/exploratory, and coping-related behaviors within a single continuous severity axis. This continuous representation better reflects the clinical spectrum from subthreshold symptoms to full disorder [6]. The high cross-cohort reproducibility observed here (ICC > 0.88) further supports MBS as a robust phenotypic readout rather than a cohort-specific construct [42].

Applying MBS across etiologically distinct stress paradigms revealed that “depression-like severity” is organized differently depending on the stress context. In CUMS, severity aligned more strongly with despair-like behavior (FST–MBS r = 0.61), whereas in CSDS it was more tightly coupled to anxiety-like exploration (OFT–MBS r = −0.74). At the same time, anhedonia-related changes were conserved across models (SPT– MBS r ≈ −0.59). Together, these results frame cross-model differences as biologically meaningful heterogeneity rather than inconsistency [5].

The stratification enabled by MBS also sharpened phenotype definition beyond unimodal criteria. In CSDS, discrepancies between MBS-based stratification and SIT-based classification (Figure 4O) indicate that single-test thresholds can miss residual impairments outside the assayed domain. MBS effectively isolates a subset of animals with broadly preserved function across multiple behavioral dimensions, which is more appropriate for mechanistic studies of resilience and for reducing within-group heterogeneity in downstream analyses. This higher-resolution stratification is also relevant for modeling partial recovery or latent vulnerability, where normalization in one dimension can coexist with persisting deficits in others [48].

Operationally, MBS simplifies behavioral assessment. Excluding TST reduces test load by 25% without loss of precision. Defined RES/SUS cohorts support targeted molecular analyses [38, 39, 40].

Several limitations should be acknowledged. While MBS provides a mathematically grounded framework for integration and stratification, the neurobiological mechanisms underlying the model-specific phenotypic architectures and resilience profiles require direct validation. In addition, although rank-based integration mitigates baseline shifts across cohorts, broader generalization across laboratories and testing conditions remains an important next step.

MBS functions as a phenotypic integrator that suppresses batch-specific artifacts while preserving biologically informative variation, enabling standardized severity quantification and high-resolution identification of resilient and susceptible subtypes. This framework supports resilience-oriented, mechanism-based investigation in preclinical depression research and may improve the interpretability and reproducibility of chronic-stress behavioral phenotyping.

## 5. Conclusion

The MBS framework establishes a novel approach for modal phenotyping in depression research by integrating individual behavioral domains into a unified severity metric. This computational approach overcomes the fragmentation in behavioral phenotyping, which has plagued conventional assessments in neuroscience and psychiatry, demonstrating exceptional cross-model reliability (ICC > 0.88 across both CUMS and CSDS models). Algorithmic stratification resolves biologically distinct endophenotypes—exemplified by stress-resilient mice preserving normative reward function despite adversity—while exposing stressor-specific phenotypic architectures. Crucially, CUMS drove despair-dominated pathology characterized by heightened FST-MBS integration (r=0.61), whereas CSDS manifested anxiety-centric dysfunction with superior OFT-MBS correlation (r=-0.74). Empirical validation helped eliminate redundant testing components (TST), reducing assessment burden without compromising accuracy. By establishing quantifiable behavioral homology across etiologically distinct models, MBS delivers a scalable phenotyping platform. Future studies leveraging this framework can dissect conserved adaptation mechanisms, accelerating the discovery of targeted interventions tailored to depression’s heterogeneous manifestations.

## Data Availability Statement

The data that support the findings of this study are available from the corresponding author upon reasonable request.

## Institutional Review Board Statement (Ethics Statement)

All animal procedures were reviewed and approved by the Animal Subjects Ethics Sub-committee of The Hong Kong Polytechnic University (approval No. 23-24/889-HTI-R-STG and No. 24-25/1083-HTI-R-STG) and were conducted in accordance with the guidelines of the Department of Health of the Hong Kong SAR.

