## supplemental figures for "Multimodal Behavior Scoring quantifies depression-like severity across chronic stress models and identifies stress-resilient mice"

**Figure S1**: Rank transformation enables robust cross-cohort integration of behavioral data in the CSDS model. (**A**) Correlation between raw and rank-transformed scores in the sucrose preference test (SPT). (**B**) Correlation between raw and rank-transformed scores in the open field test (OFT). (**C**) Correlation between raw and rank-transformed scores in the forced swim test (FST). (**D**) Rank-based trajectories of individual behavioral metrics (SPT, OFT, FST) sorted by ascending MBS scores in CSDS control group. (**E**) Rank-based trajectories of individual behavioral metrics (SPT, OFT, FST) sorted by ascending MBS scores in CSDS stressors group. (**F**) Ordered bar chart of MBS in CSDS control group. (**G**) Ordered bar chart of MBS in CSDS stressors group.

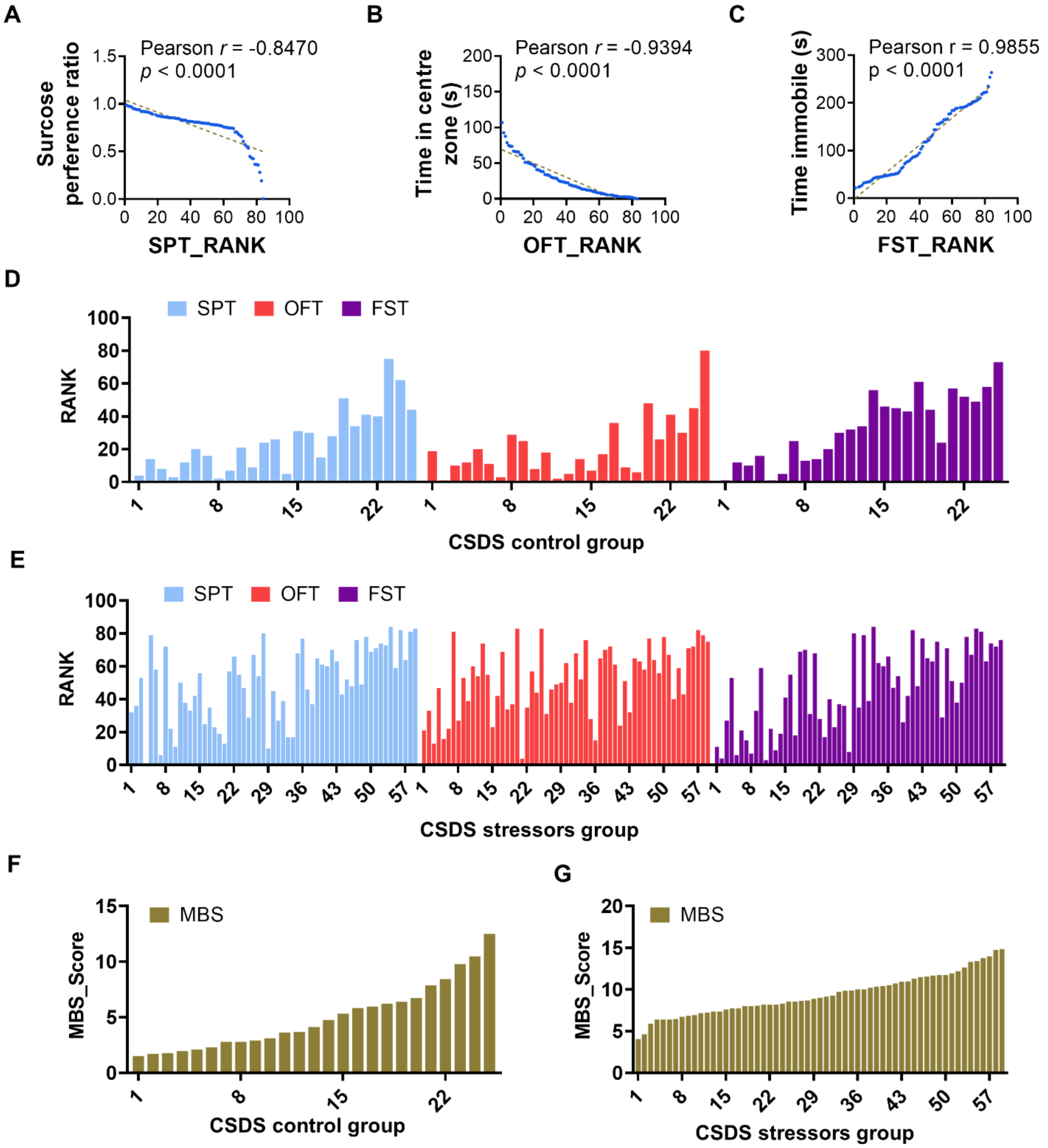

**Figure S2**: (**A**) Mouse-specific rank-based trajectories of individual behavioral metrics (SPT, OFT, FST) sorted by ascending MBS scores in the CUMS control group. (**B**) Mouse-specific rank-based trajectories of individual behavioral metrics (SPT, OFT, FST) sorted by ascending MBS scores in the CUMS control group.

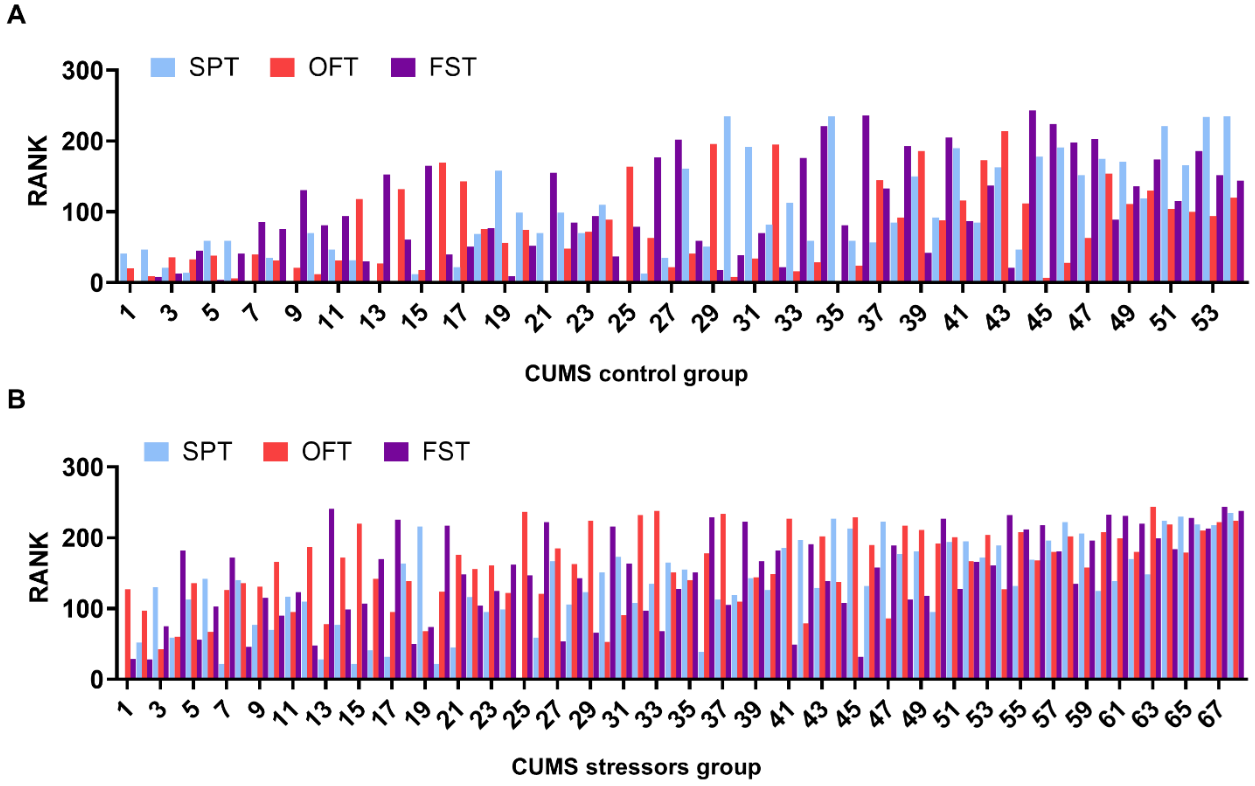

**Figure S3**: (**A**) Mouse-specific rank-based trajectories of individual behavioral metrics (SPT, OFT, FST) sorted by ascending MBS scores in the CSDS control group. (**B**) Mouse-specific rank-based trajectories of individual behavioral metrics (SPT, OFT, FST) sorted by ascending MBS scores in the CSDS control group.

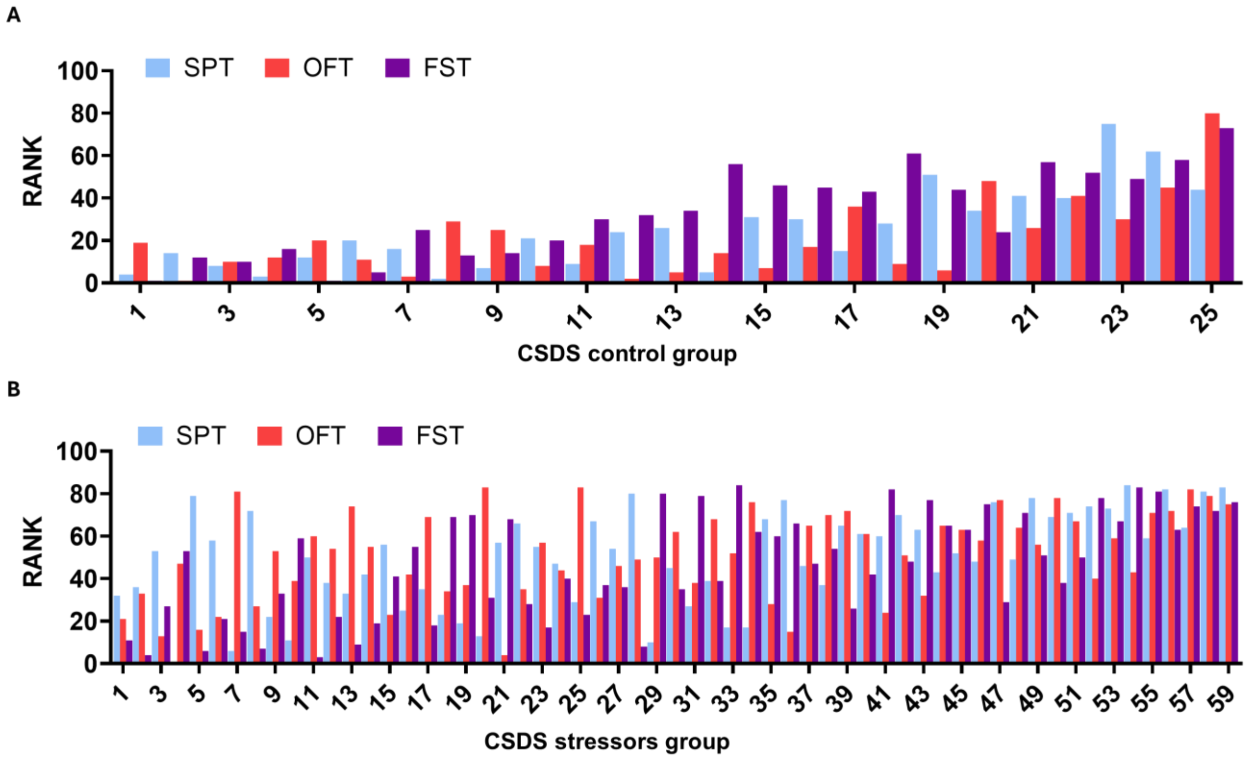

**Figure S4**: MBS scores and three behavioral indicators of two cohorts of mice in the CUMS model.

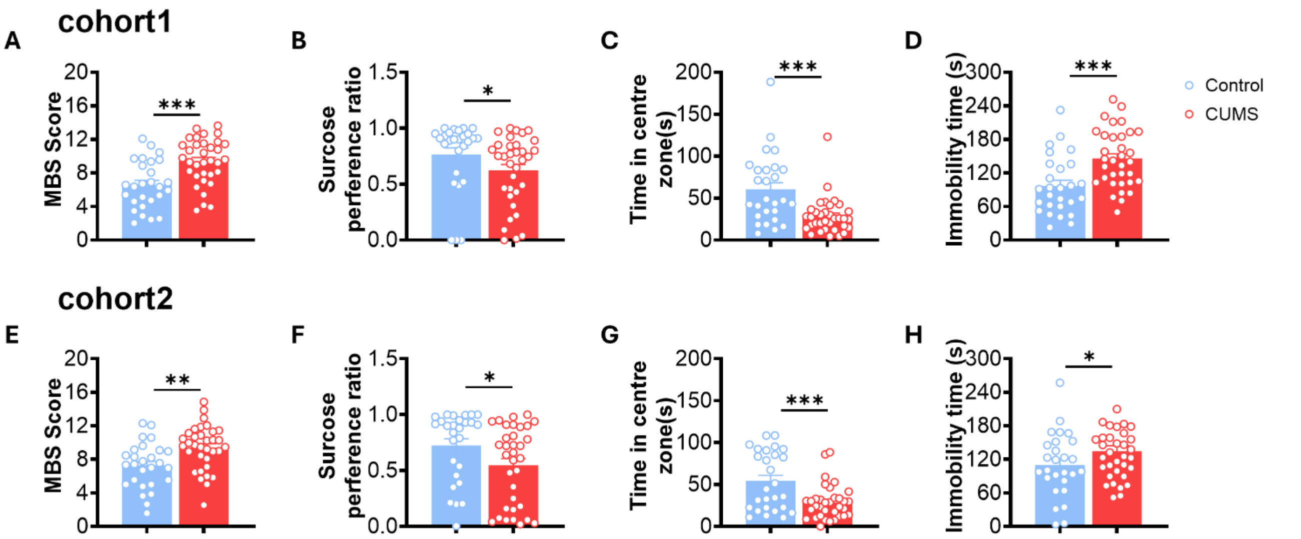


**Figure S5**: MBS scores and three behavioral indicators of two cohorts of mice in the CSDS model.

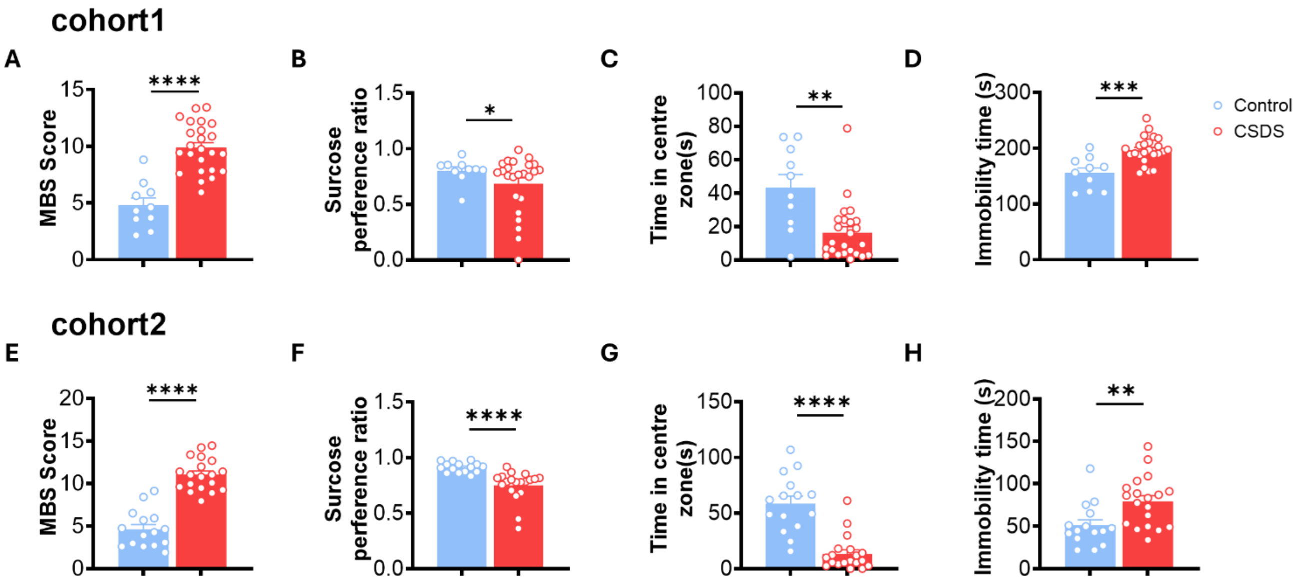
